# Prevention and Treatment of SHIVAD8 Infection in Rhesus Macaques by a Potent D-peptide HIV Entry Inhibitor

**DOI:** 10.1101/2020.04.14.041764

**Authors:** Y. Nishimura, J.N. Francis, O. Donau, E. Jesteadt, R. Sadjadpour, A.R. Smith, M.S. Seaman, B.D. Welch, M.A. Martin, M.S. Kay

## Abstract

Cholesterol-PIE12-trimer (CPT31) is a potent D-peptide HIV entry inhibitor that targets the highly conserved gp41 N-peptide pocket region. CPT31 exhibited strong inhibitory breadth against diverse panels of primary virus isolates. In a SHIV macaque model, CPT31 prevented infection from a single high-dose rectal challenge. In chronically infected animals, CPT31 monotherapy rapidly reduced viral load by ~2 logs before rebound occurred due to the emergence of drug resistance. In chronically infected animals with viremia initially controlled by combination antiretroviral therapy (cART), CPT31 monotherapy prevented viral rebound after discontinuation of cART. These data establish CPT31 as a promising new candidate for HIV prevention and treatment.

## Introduction

Combination antiretroviral therapy (cART) has greatly improved the length and quality of life of HIV infected individuals with access to treatment and has reduced HIV transmission from treated patients (Cohen et al., 2016; Egger et al., 2002; Hogg et al., 2002; Palella et al., 1998). Despite continuing improvements, cART remains costly, has toxic side effects, and requires daily administration. Additionally, the high mutation rate of HIV-1 results in the rapid development of resistance to all FDA-approved antiretroviral drugs (Johnson et al., 2010). As a result, there is an ongoing need for novel, cost-effective therapeutic options with unique mechanisms of action and the potential for extended dosing.

We previously described the development and characterization of a D-peptide entry inhibitor, cholesterol-PIE12-trimer (CPT31), which exhibits low-picomolar (pM) potency against a Tier 2 HIV-1 strain (HIV-1_JRFL_) and has a favorable pharmacokinetic (PK) profile in non-human primates (Francis et al., 2012; Redman et al., 2018). PIE12 is a 16-residue D-peptide (composed of D-amino acids) that binds to the highly conserved gp41 trimer “pocket” region, which plays a key role in mediating viral membrane fusion (Chan et al., 1998). A trimeric version of PIE12, connected via flexible PEG linkers (PIE12-trimer), binds with high avidity to the three gp41 pockets in the HIV-1 envelope trimer, and was previously shown to have a very high affinity for trimeric gp41 (Welch et al., 2010). In tissue culture virus-passaging studies, resistance to PIE12-trimer was mediated by a gp41 pocket region mutation (typically Q577R) (Smith et al., 2019; Welch et al., 2010). Conjugation of PIE12-trimer to cholesterol (to produce CPT31) localizes the inhibitor to the membrane sites of viral entry, further enhancing potency (Francis et al., 2012; Redman et al., 2018).

We have previously used the rhesus macaque and R5-tropic simian-human immunodeficiency virus (SHIV) AD8 system as a surrogate for human HIV-1 infections because it exhibits multiple clinical features observed in the human disease, following either intravenous or intrarectal inoculation (Gautam et al., 2012; Nishimura et al., 2010; Shingai et al., 2012). Unlike most other SHIVs, SHIVAD8 generates sustained levels of viremia, resulting in the slow and continuous loss of CD4^+^ T lymphocytes, the development of associated opportunistic infections and lymphomas, weight loss, and death within 2 to 4 years. Virus replication in animals challenged with an infectious molecular clone derived from SHIVAD8, designated SHIVAD8-EO, can be transiently controlled with broadly neutralizing antibody (bNAb) monotherapy, but resistant viral variants invariably and rapidly emerge (Shingai et al., 2013). In this study, we report that CPT31 is efficacious as both a preventative and a therapeutic agent against SHIVAD8-EO infections of rhesus macaques.

## Results

### CPT31 inhibitory breadth

The inhibitory breadth of CPT31 was initially characterized using the CAVD 118-strain pseudovirion panel (Huang et al., 2016; Seaman et al., 2010). This group of pseudovirions includes strains from clades A, B, C, D, and G, as well as circulating recombinant forms AC, ACD, AE, AG, BC, and CD. All 118 HIV-1 pseudovirions were fully inhibited by CPT31, with IC_50_s varying from <1 to 490 pM (average 50 pM) (Fig. 1A and Table S1). None of the strains in this HIV-1 Env panel carried the Q577R gp41 resistance mutation.

**Figure 1:**
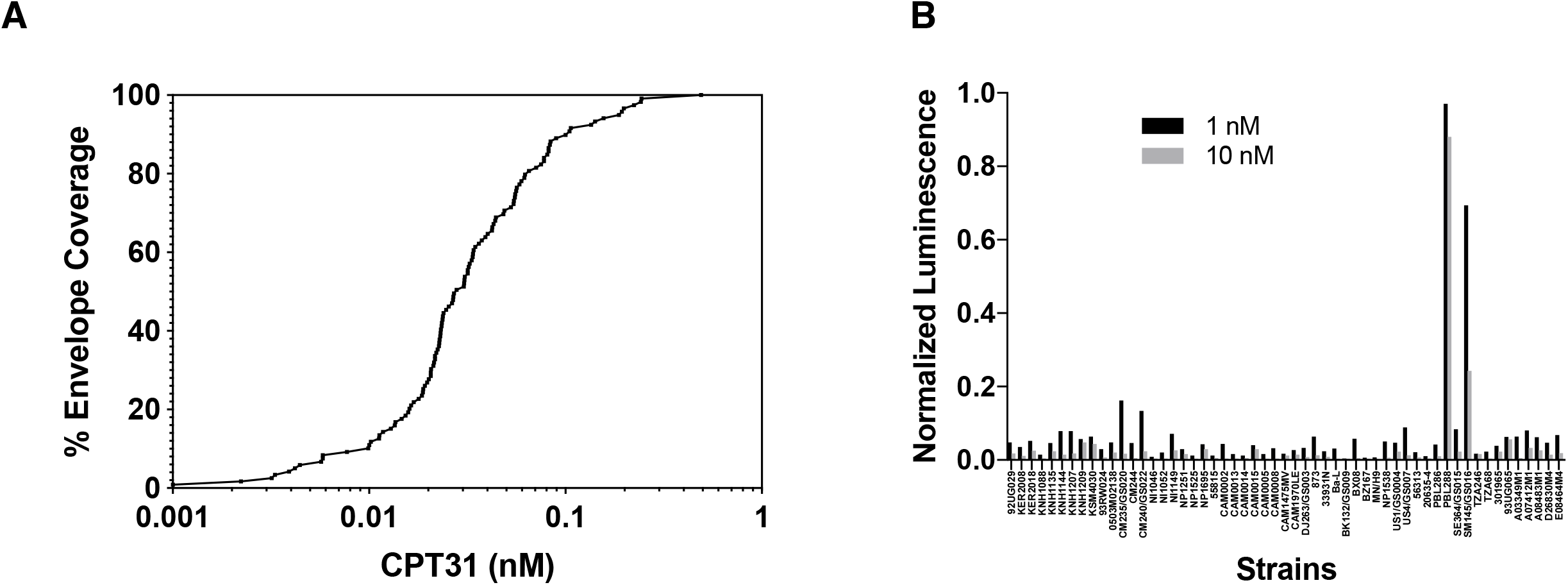
The inhibitory breadth of CPT31 against panels of HIV-1 isolates. A) Percentage of strains inhibited at least 50% (IC_50_) at the indicated concentrations using the CAVD 118-strain HIV-1 pseudovirion panel. B) Normalized luciferase values from 60-strain international panel of replication-competent HIV-1 strains obtained from the NIH AIDS Reagent Program and used at 1 and 10 nM CPT31.

The inhibitory breadth of CPT31 was also evaluated using an international panel of 60 primary replication-competent isolates (#8129) obtained from the NIH AIDS Reagent Program, and the infectivity of these viruses was assessed in TZM-bl cells. This panel includes 60 isolates from Clades A, B, C, D, and circulating recombinant forms AE and AG. Three of the 60 panel viruses failed to produce luciferase levels above background and were excluded from further study. Of the 57 remaining HIV-1 strains, 55 (96%) were >90% inhibited at 10 nM CPT31, and 53 (93%) were >90% inhibited at 1 nM CPT31 (Fig. 1B and Table S2). The two HIV-1 strains (PBL288 and SM145) exhibiting weak inhibition at 10 nM carry the previously described CPT31 gp41 resistance mutation (Q577R) (Welch et al., 2010). The inhibitory CPT31 breadth exhibited against HIV-1 isolates from both panels, combined with its favorable PK profile in NHPs (Redman et al., 2018), encouraged us to assess the efficacy of CPT31 in controlling SHIVAD8-EO infections of non-human primates.

### Prevention of virus acquisition by CPT31 in rhesus macaques

Results from multiple studies have shown intrarectal (IR) inoculation of 1000 TCID_50_ of the uncloned SHIVAD8 swarm virus stock or its molecularly cloned SHIVAD8-EO derivative (Gautam et al., 2012; Shingai et al., 2012) results in the successful establishment of infection in rhesus macaques. Representative virus replication profiles of SHIV AD8-EO, as measured by the levels of plasma viral RNA, are shown in Fig. 2A ((Shingai et al., 2012) and unpublished results). To assess the capacity of CPT31 to block the establishment of SHIVAD8-EO infections, 4 monkeys were initially treated with intramuscular (IM) CPT31 injections (3 mg/kg/day) for 10 days. On day 3 of therapy, the animals were challenged with 1000 TCID_50_ SHIVAD8-EO by the IR route. As shown in Fig. 2B, all four macaques remained uninfected for 21 weeks, as measured by RT-PCR analyses of sequential plasma samples. To verify that SHIVAD8-EO acquisition had not occurred in the protected monkeys, whole blood from these four animals was pooled and inoculated into a single naïve rhesus macaque. Specifically, PBMCs and plasma, isolated from 20 ml of blood from the each of the four protected monkeys at week 18 post-challenge, were pooled and infused intravenously into a naive animal. This macaque failed to become infected, as monitored by multiple RT-PCR assays of plasma during a 40-week observation period.

**Figure 2:**
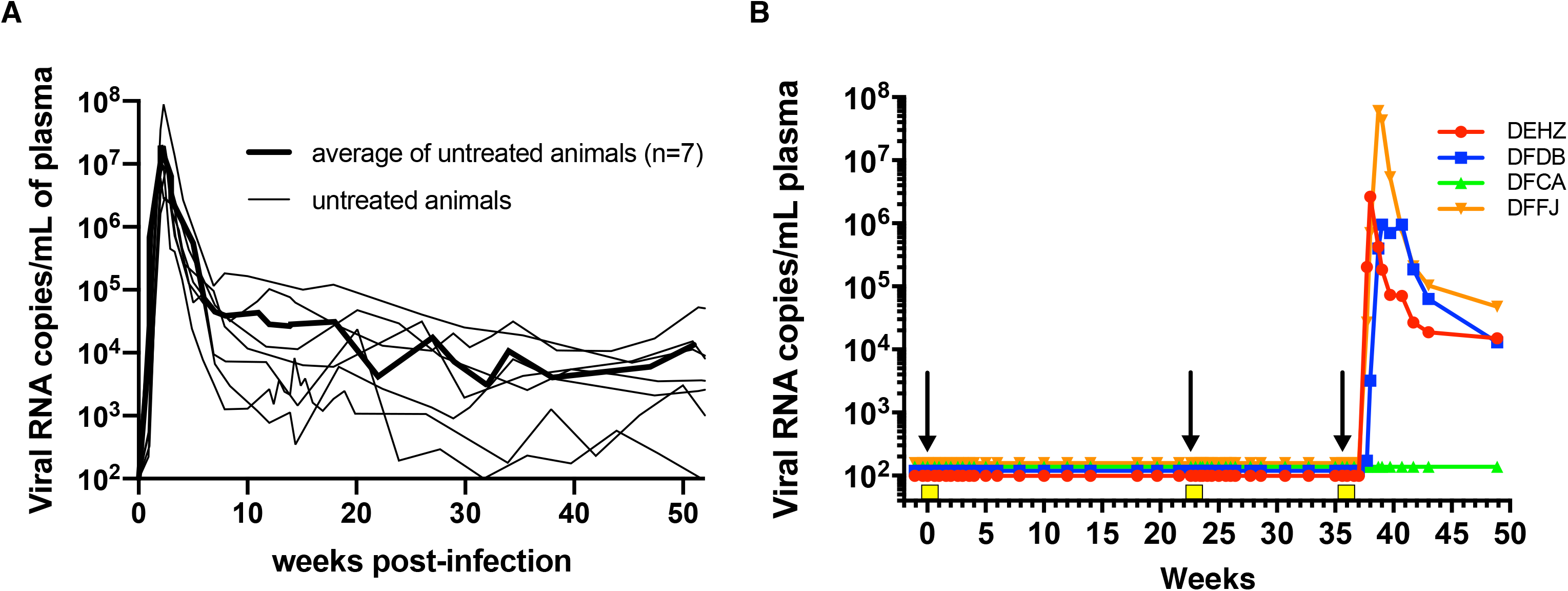
CPT31 is able to block virus acquisition in rhesus macaques. A) Levels of plasma viremia in seven previously described control rhesus macaques following intrarectal inoculation with 1000 TCID_50_ of SHIVAD8-EO (Shingai et al., 2012). The average viral loads in these seven animals is indicated by the bold line. B) Four monkeys were inoculated intrarectally with 1000 TCID_50_ of SHIVAD8-EO during weeks 0, 22 and 35 and intramuscularly administered 3.0, 0.5, or 0.125 mg/kg CPT31 daily, from day –3 to day +7, at each time of virus challenge. Arrows indicate the times of viral challenges; the 10 days of the three CPT31 treatments are shown as yellow boxes.

The same 4 animals were then administered a second 10-day course of a lower CPT31 dose (0.5 mg/kg/day), initiated at week 22 post-challenge, and intrarectally re-challenged with 1000 TCID_50_ of SHIV AD8-EO 3 days later. The administered CPT31 again prevented virus acquisition for an additional 13 weeks (Fig. 2B). Finally, the same 4 animals were treated with an even lower dose of inhibitor (0.125 mg/kg/day) for 10 days beginning at week 35 and inoculated IR a third time with 1000 TCID_50_ of SHIV AD8-EO 3 days later. In this case, plasma viremia was measured beginning at week 37 in 3 of the 4 monkeys. One animal remained uninfected at week 49 when the experiment was terminated. SHIV RNA was isolated and PCR-amplified from the plasma of the 3 infected animals to ascertain whether resistant variants had emerged. Viral RNA from all three animals were >99% WT at position Q577 (>100,000 reads each) indicating that insufficient CPT31 dosing, not drug resistance, was responsible for the viremia observed in these monkeys. Taken together, these data suggest a minimum protective dose for CPT31 in the high-dose rectal challenge SHIV AD8-EO/rhesus macaque model is between 0.5 and 0.125 mg/kg/day.

CPT31 plasma levels were measured in all 4 animals during the “dose-de-escalation” experiments, using an LC-MS bioanalytical assay (Fig. S1). For the 3, 0.5, and 0.125 mg/kg/day treatments, the average drug levels for samples taken immediately prior to each day’s injection were 347, 94, and 25.5 nM, respectively. The SHIVAD8-EO protective CPT31 plasma concentration was therefore between 94 and 25.5 nM. The decay of drug levels after discontinuation of each dosing was also monitored (Fig. S1). CPT31 levels fell below the detection limit of 1 nM in 7 weeks, 11 days, and 4 days for the 3, 0.5, and 0.125 mg/kg/day dosing regimens, respectively. These drug levels are similar to those predicted by the subcutaneous pharmacokinetic parameters previously reported in cynomolgus monkeys (Redman et al., 2018).

### CPT31 treatment during chronic SHIV AD8-EO infections of rhesus macaques

Chronic virus infections were established in 3 rhesus monkeys following the IR inoculation of 1000 TCID_50_ of SHIV AD8-EO. Peak levels of virus production in the 10^7^ to 10^8^ viral RNA copies/ml plasma range were measured at week 2 post-infection (PI) and set points of plasma viremia in the range of 10^4.1^ to 10^5.6^ viral RNA copies/ml were established between weeks 10 to 14 in the 3 animals (Fig. 3). A 30-day course of CPT31 monotherapy (3 mg/kg/day/IM) was initiated at week 14 PI. All three monkeys experienced a rapid ~2-log decline in viral load within 1 to 2 weeks of treatment initiation indicating that CPT31 was able to potently suppress virus replication *in vivo.* However, virus rebound occurred in all three animals a week or two later. Because drug levels monitored in the 4 monkeys indicated that plasma CPT31 concentrations remained at the 200 nM level during the 30-day period of inhibitor administration (Fig. S2), the emergence of CPT31-resistant virus seemed likely.

**Figure 3:**
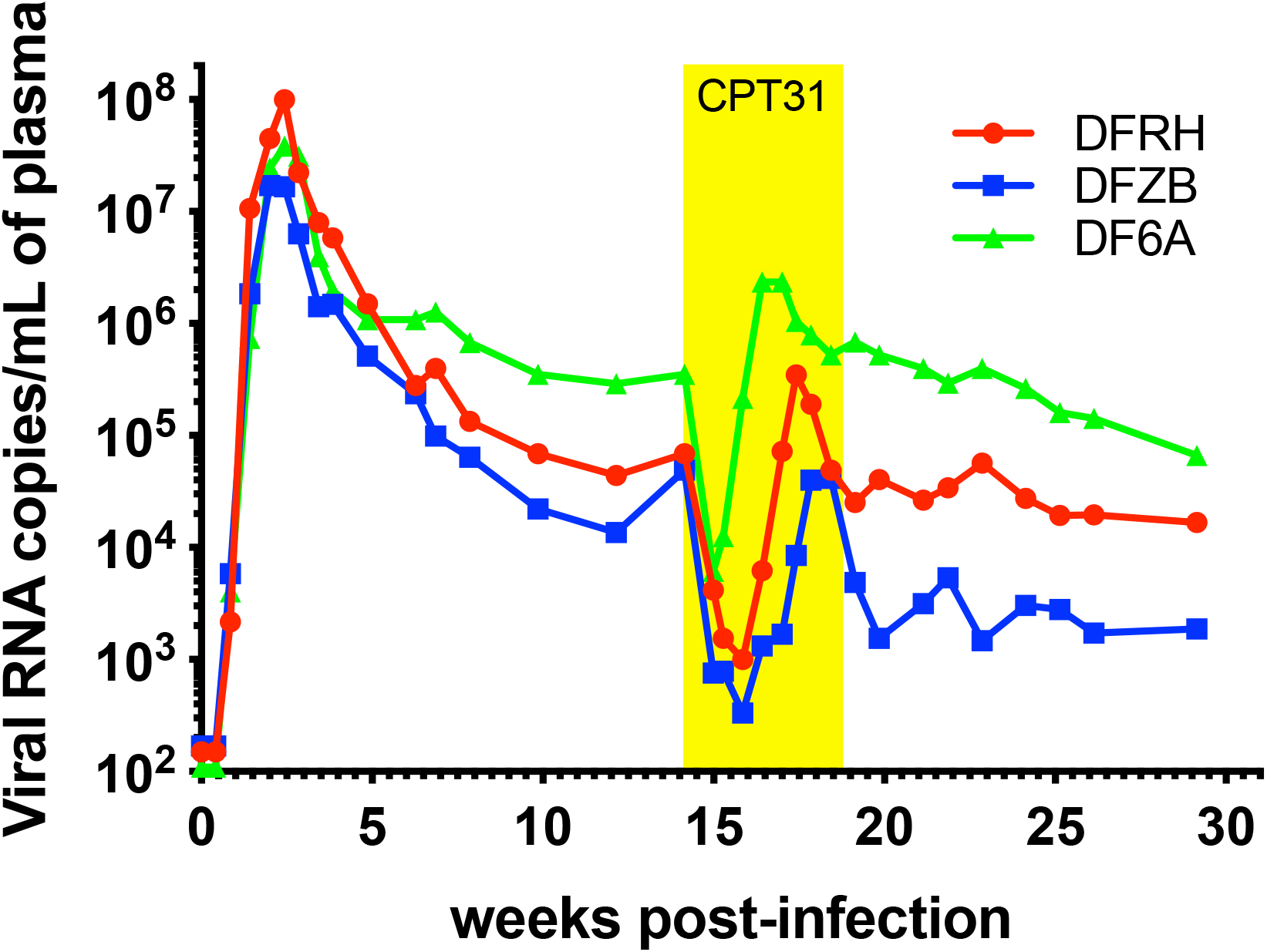
CPT31 monotherapy of chronically SHIVAD8-EO infected rhesus macaques. Three chronically SHIVAD8-EO infected animals were treated for 4 weeks with 3 mg/kg/day CPT31, beginning at week 14 PI.

To ascertain whether resistance to CPT31 had, in fact, occurred during anti-viral monotherapy, plasma was collected from the 3 treated macaques at weeks 18 and 30 PI, viral RNA was RT-PCR amplified, and *env* genes sequenced. As shown in Fig. 4A, virtually all of the virus circulating in the 3 animals at week 18 PI, when SHIVAD8-EO had rebounded during CPT31 treatment, carried the Q577R CPT31 resistance substitution previously reported to arise in tissue culture passaging studies (Smith et al., 2019; Welch et al., 2010). Interestingly, 14 of 20 *env* genes cloned from macaque DF6A at week 18 PI had a downstream S668G change, located in the membrane-proximal external region (MPER) of gp41, in addition to the Q577R substitution.

**Figure 4:**
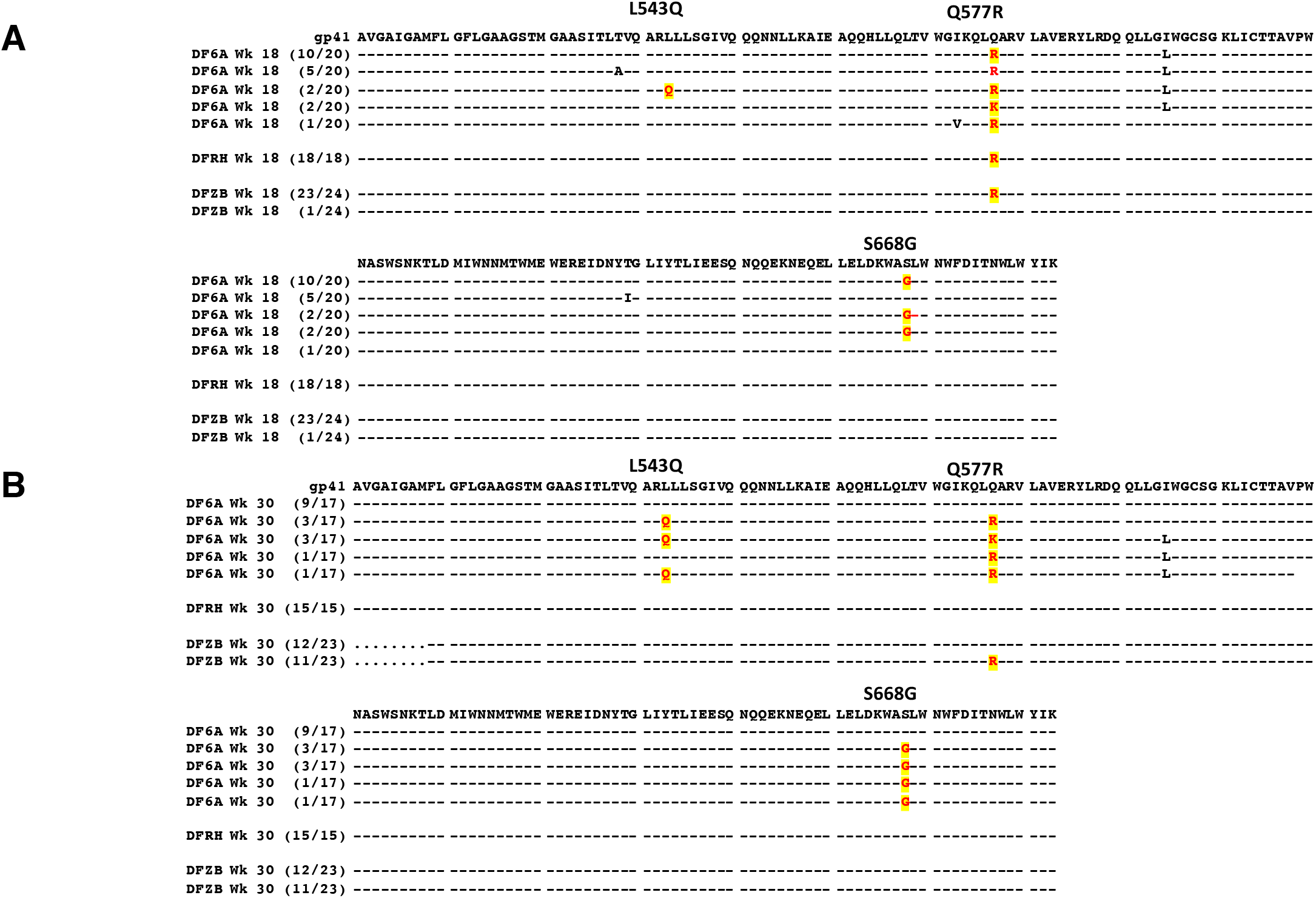
Sequence analyses of gp41 gene segments in SHIVAD8-EO chronically infected macaques treated with CPT31 monotherapy. Viral RNA was amplified by RT-PCR from plasma collected during (week 18 [A]) and following (week 30 [B]) CPT31 treatment. Common amino acid changes at positions 543, 577 and 668 of gp41 in the three animals and their frequencies are highlighted. Residues 511-683 are shown.

At week 30 PI, the Q577R change had completely reverted to wild type (WT) in animal DFRH (Fig. 4B). In contrast, 11 of 23 cloned *env* genes amplified from the plasma of macaque DFZB at week 30 PI retained the Q577R substitution. The SHIVAD8-EO circulating in monkey DF6A at week 30 PI was genetically more complex: 8 of 17 of the amplified *env* genes retained the Q577R substitution as well as the S668G change; 7 of these same 8 *env* genes had also acquired a new L543Q change (Fig. 4B), located in the N-heptad repeat region of gp41.

### Biological properties of SHIV AD8-EO CPT31-resistant variants

Molecularly cloned derivatives of SHIVAD8-EO, carrying the *env* gene Q577R (AD8-577R) or Q577R plus L543Q (AD8-577R/543Q) substitutions, were constructed and used to generate virus stocks by transfecting 293T cells. The resulting supernatants were used to infect rhesus PBMCs to generate infectious virus stocks. The sensitivity of WT SHIVAD8-EO, plus the AD8-577R and AD8-577R/543Q resistant variants, to CPT31 was evaluated using *in vitro* infectivity assays with 4-fold serial dilutions of the D-peptide, as described in Methods. The autoradiograms in Fig. 5A indicate that infection of WT SHIV AD8-EO was 50% blocked at the 4^−4^ dilution (1.6 nM CPT31), whereas the AD8-577R and AD8-577R/543Q SHIV variants, which emerged during the D-peptide treatment *in vivo*, were both resistant at the highest concentration (100 nM) tested.

**Figure 5:**
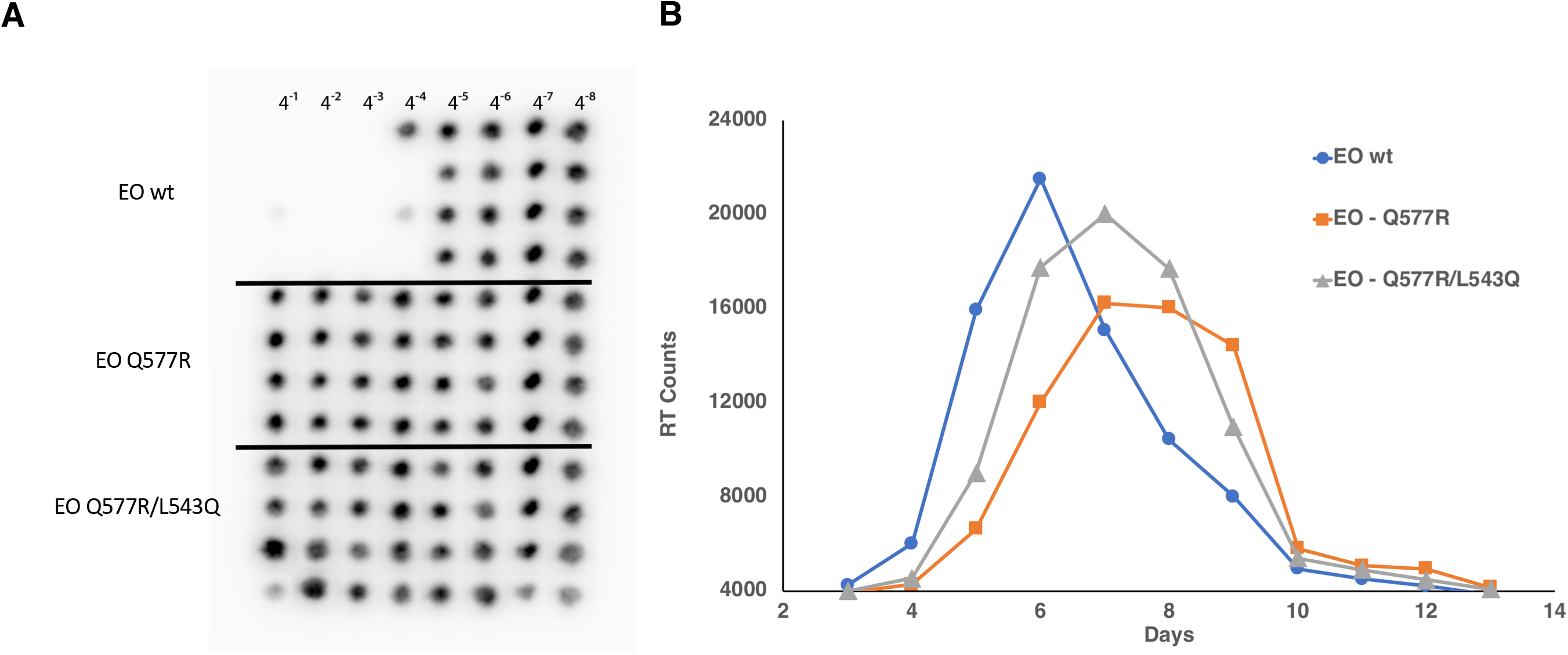
Characterization of SHIVAD8-EO CPT31-resistant variants. A) Endpoint titrations of CPT31 inhibitory activities were assessed against wild type or resistant SHIVAD8-EO variants in rhesus PBMC. The presence or absence of progeny virus production, treated with the serially diluted CPT31 (four-fold), was measured autoradiographically by ^32^P-reverse transcriptase assays performed on aliquots of the culture supernatant from day 21 of infection. The black spots in the autoradiograms indicate the presence of virion-associated reverse transcriptase activity (i.e., no blocking of virus replication). B) Replication of wild type and SHIVAD8-EO variants in rhesus monkey PBMC. Virus stocks prepared in rhesus PBMC were used to infect rhesus PBMC (MOI=0.002). Virus replication was assessed by RT activity released into the culture medium.

The infectivities of WT SHIVAD8-EO, AD8-577R, and AD8-577R/543Q SHIV variants were next assessed using an MOI of 0.002 of each virus stock spinoculated onto 1×10^6^ rhesus PBMC cultures. As shown in Fig. 5B, the replication of the AD8-577R SHIV variant was delayed by 1 to 2 days compared to WT SHIVAD8-EO, and the AD8-577R/543Q SHIV variant was delayed by 1 day. Taken together, the amino acid substitutions conferring CPT31 resistance had modest effects on SHIV AD8-EO replication fitness *in vitro*. This result was consistent with the robust SHIV replication kinetics observed in all infected macaques following virus rebound (Fig. 3), including monkey DF6A, in which nearly half of the circulating virus population was carrying the Q577R and the L543Q variants at week 30 PI.

### CPT31 monotherapy controls virus replication in chronically SHIVAD8-EO infected macaques when administered following virus suppression conferred by prior conventional combination anti-retroviral treatment

The transient effect of CPT31 monotherapy administered to chronically SHIVAD8-EO infected macaques with set point levels of viremia in the 4 to 5 log_10_ range shown in Fig. 3 was not unexpected given the high levels of sustained production of progeny virus in these animals. We wondered whether potentially potent properties of the D-peptide *in vivo* might be revealed in the context of an established chronic infection if virus replication was initially suppressed with conventional combination anti-retroviral therapy (cART), and the efficacy of CPT31 monotherapy assessed following cessation of preceding and partially overlapping ART. This possibility was evaluated by first treating four chronically SHIVAD8-EO macaques with a 5-week course of cART (emtricitabine/tenofovir/raltegravir) starting at week 19 PI. As shown in Fig. 6, this therapy resulted in the rapid decline of plasma viremia to background levels in all 4 monkeys, demonstrating that a conventional cART regimen effectively suppresses SHIVAD8-EO replication. Virus was allowed to rebound for 15 weeks following cessation of cART treatment at week 24 PI. At week 41 PI, a 6-week course of cART was initiated, but in this case, CPT31 (3 mg/kg/day) was started in the last week of cART treatment (at week 46 PI) and continued for an additional 13 weeks (to week 59 PI). Fig. 6 shows that virus replication was suppressed to background levels during the period of CPT31 monotherapy, and rapid rebound of plasma viremia occurred within 3 weeks in all 4 treated animals following cessation of daily D-peptide administration. The measurement of plasma CPT31 concentrations during and after the 12 weeks of inhibitor monotherapy are shown in Fig. S3, and viral rebound occurs as expected with the drop in CPT31 to below therapeutic levels. Sequencing of the *env* genes in viruses emerging, between weeks 62 to 64, following the discontinuation of D-peptide treatment, revealed that: 1) macaques DFFF, DFNH and DFNW carried none of the previously observed CPT31 resistance substitutions and 2) monkey DFTV had 9/21 *env* clones with the Q577R change, 7 of which, also carried the 543Q substitution (Fig. S4).

**Figure 6:**
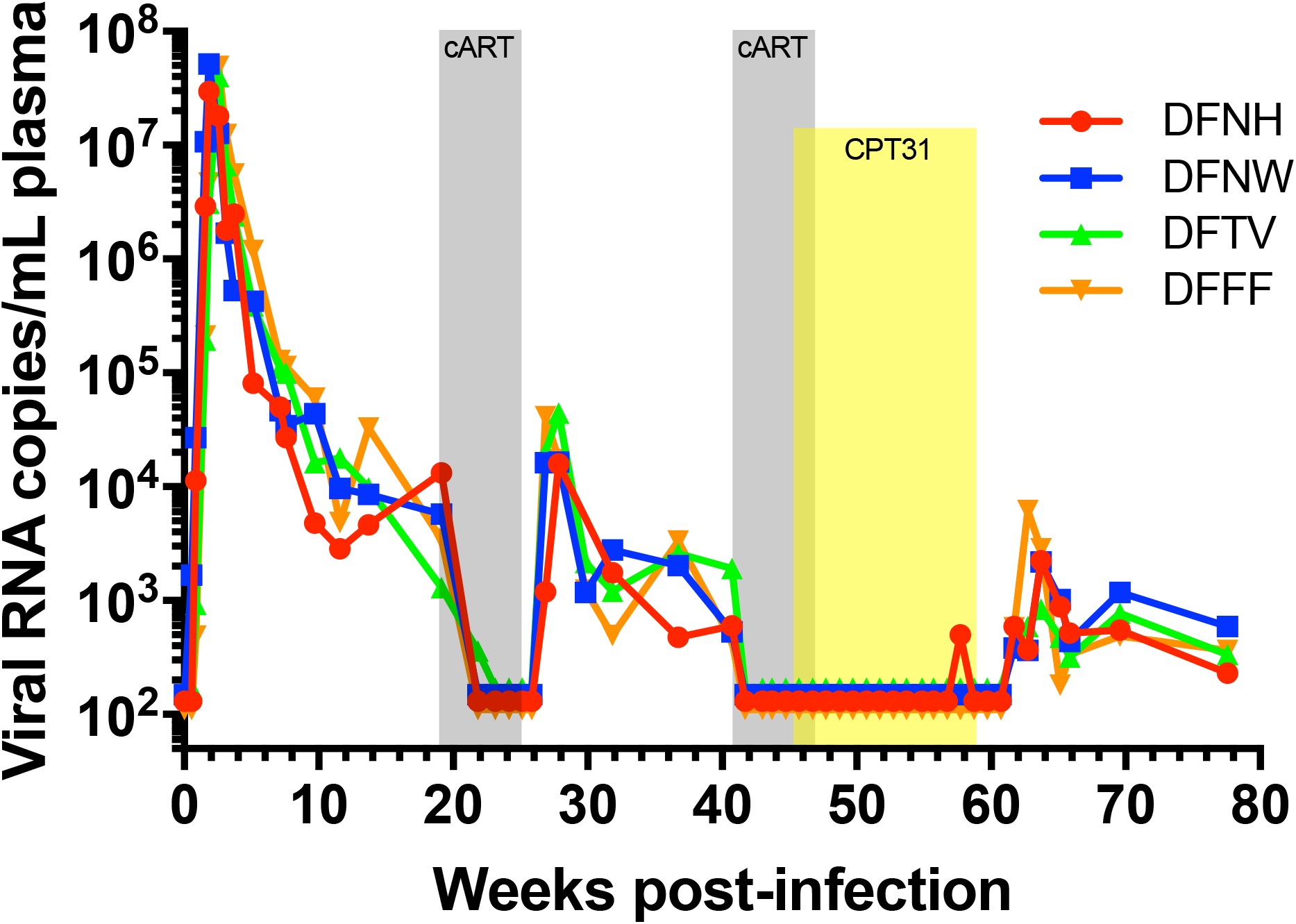
CPT31 monotherapy is able to control viremia in chronically virus infected macaques if SHIVAD8-EO replication is first suppressed by combination anti-retroviral treatment. Four monkeys were treated with cART (emtricitabine/tenofovir and raltegravir) starting at week 19 for 5 weeks (gray) and at week 41 PI for an additional 6 weeks. CPT31 (3 mg/kg/day) was added in the last week of cART (at week 46 PI) and continued as monotherapy for an additional 12 weeks (yellow).

## Discussion

In this study, we used the SHIV/macaque model to investigate the efficacy of CPT31 in both therapeutic and preventative contexts. CPT31 conferred complete protection against a high-dose rectal challenge when dosed at 0.5 mg/kg/day and was partially protective at 0.125 mg/kg/day. Allometric scaling between macaques and humans predicts a 3-fold reduction in the drug dosing required to obtain comparable serum levels (FDA, 2005). Therefore, we hypothesize that ≪0.125 mg/kg/day dosing would protect against more physiologic (lower-dose) mucosal transmission in humans. Since CPT31 is not a component of existing cART regimens, development of CPT31 resistance as a consequence of previous participation in PrEP (e.g., in patients with undocumented infections) would be unlikely to affect cART treatment options.

The broad efficacy of CPT31 against diverse panels of HIV-1 isolates shown in Fig. 1 suggests it will be effective against a wide range of viral strains. The protective breadth of CPT31 indicates potent inhibition of all strains tested across diverse clades except for those bearing the gp41 Q577R resistance mutation. We anticipate that CPT31 will be ineffective against Group O strains (Welch et al., 2010), which have a high prevalence of the Q577R primary resistance mutation (>99% Q577R in Group O strains) vs ~2% Q577R in group M (Los Alamos National Lab HIV Sequence Database). The impact of this limitation is expected to be minimal given the relatively low prevalence of Group O infections worldwide (Villabona-Arenas et al., 2015).

CPT31 monotherapy of an established infection rapidly inhibits viral replication, but as expected for a highly active compound in the context of uncontrolled viremia, induces drug resistance. The primary resistance mutation observed in macaques (Q577R) is in agreement with that reported in prior *in vitro* studies (Welch et al., 2010). The additional gp41 mutations observed *in vivo* (in the MPER (S668G) and N-terminal heptad repeat (543Q)) are of unknown significance and were not observed in the context of *in vitro* resistance to PIE12-trimer (Smith et al., 2019). These mutations have also not been reported in the context of resistance to other HIV entry inhibitors. Their effect on CPT31 resistance is likely to be indirect (e.g., affecting fusion kinetics, gp41 pocket accessibility, and/or compensating for fitness defects associated with the Q577R substitution). We also monitored whether CPT31 monotherapy alone could maintain the previous ART-mediated suppression of virus replication in chronically infected animals. Such CPT31 monotherapy potently controlled viral replication for an additional 12 weeks, at which point CPT31 was discontinued.

Overall, the preclinical data presented support CPT31 as a strong candidate for both PrEP and as a component of combination therapy against a wide variety of HIV isolates. Our current potency and PK results suggest that CPT31 would be suitable for depot formulation and, ideally, could be used as a long-acting injectable for monthly or less frequent dosing. Studies are underway to formulate CPT31 in a depot that could be used in such a context. Additionally, this formulation could be paired with other long-acting drugs, such as nanocrystals of cabotegravir (GSK744, GlaxoSmithKline) and/or rilpivirine (TMC278, Tibotec) (Orkin et al., 2020; Swindells et al., 2020), ibalizumab (Emu et al., 2018), or promising antibodies and small molecule inhibitors (Merck’s islatravir/MK-8591 and Gilead’s GS-6207) in clinical development (Coelho et al., 2019; Schurmann et al., 2020). To fully evaluate CPT31’s therapeutic potential, it will be important to evaluate its efficacy when administered with other antiretrovirals to identify optimal combinations. The FDA recently cleared CPT31’s Investigational New Drug (IND) application, and a first-in-human trial is planned for 2020.

## Methods

### Animal Experiments

Eleven male and female RMs (*Macaca mulatta*) of Indian genetic origin ranging from 2 to 4 years of age were maintained in accordance with the guidelines of the Committee on Care and Use of Laboratory Animals (Department of Health and Human Services, Bethesda, MD, 1985) and were housed in a biosafety level 2 facility; biosafety level 3 practices were followed. Phlebotomies, intrarectal (IR) virus inoculations, intramusccular (IM) drug administration, and sample collections were performed as previously described (Endo et al., 2000; Gautam et al., 2012; Nishimura et al., 2017). All animals were negative for the major histocompatibility complex class I *Mamu-A*\**01*, *Mamu-B*\**08*, and *Mamu-B*\**17* alleles.

The origin and preparation of the tissue-culture-derived SHIV_AD8-EO_ stock have been previously described (Shingai et al., 2012). Animals were challenged with 1,000 TCID_50_ of SHIVAD8-EO intrarectally, as previously described (Shingai et al., 2014).

CPT31 was formulated at 10 mg/mL in PBS (50 mM sodium phosphate, 150 mM NaCl, pH 7.4) and administered IM on a mg/kg basis. Four animals were treated with a three-drug ART regimen comprising two nucleoside reverse transcriptase (RT) inhibitors (tenofovir (PMPA) and emtricitabine (FTC)) and one integrase inhibitor (raltegravir (RAL)), starting at week 19 for 5 weeks and starting at week 46 PI for 6 weeks. PMPA and FTC were administered intramuscularly once a day at dosages of 20 mg/kg and 40 mg/kg, respectively. RAL was administered orally (mixed with food) at a dosage of 200 mg twice a day.

### Quantitation of plasma viral RNA levels

Viral RNA levels in plasma were determined by qRT–PCR (ABI Prism 7900HT sequence detection system; Applied Biosystems) as previously described (Endo et al., 2000).

### Lymphocyte immunophenotyping

EDTA-treated blood samples were stained for flow cytometric analysis for lymphocyte immunophenotyping as previously described (Nishimura et al., 2010).

### Virus Sequencing

Viral RNA was isolated from macaque plasma using the QIAmp Viral RNA Isolation kit (QIAGEN) according to manufacturer protocol and converted to cDNA using the Superscript III or Superscript IV First Strand cDNA synthesis kit (ThermoFisher). 2 μL of cDNA was subject to PCR amplification for the D-peptide binding region using Platinum PCR Supermix HiFi (Thermo) 5’-TGTATGCCCCTCCCATCAGA-3’ (forward) and 5’-CAAGCGGTGGTAGCTGAAGA-3’ (reverse) primers (Integrated DNA Technologies) in a 50 μL reaction. Initial denaturation was carried out at 94°C for 2 minutes, followed by 32 cycles of 20 seconds at 94°C, 30 seconds at 55°C and 2 minutes at 68°C, with a final extension for 7 minutes at 68°C. PCR products were ligated into PCR4-TOPO-TA vectors using the TOPO-TA Cloning Kit (ThermoFisher) according to manufacturer instructions. 2 μL of each ligation reaction was transformed into TOP10 Chemically competent *E. Coli* (ThermoFisher) according to manufacturer instructions. Transformants were plated on LB agar plates containing 100 μg/mL ampicillin and were allowed to incubate overnight at 37°C. Single colonies were inoculated into 3 mL cultures containing LB medium with 100 μg/mL ampicillin and were allowed to incubate overnight at 37°C with 250 rpm shaking. 2-3 mL of each bacterial culture was pelleted and plasmid DNA extracted and purified using the QIAPrep SpinPrep miniprep kit (QIAGEN) according to manufacturer instructions. 2 μg of each clone was sent to the NIAID LMM-CORE (Twinbrook, MD) for Sanger sequencing. Sequences were aligned using MacVector sequence analysis software (MacVector Inc.).

Next-generation sequencing was conducted by using viral RNA isolated from three monkey plasma samples (DEHZ 5/21/2016, DFFJ 6/1/2016, and DFDB 6/3/2016). The isolated RNA samples, using the E.Z.N.A Viral RNA kit (Omega), were reverse transcribed using SuperScript IV (ThermoFisher) and then amplified using nested PCR with primers specific to SHIV_AD8_ Env as well as individual barcodes for deep sequencing. The amplified samples were analyzed at the DNA sequencing core at the University of Utah using the Ion Torrent PGM Next-Generation Sequencing Platform.

### Construction of CPT31 resistant viruses

The Q577R gp41 change was introduced into pSHIV-AD8-EO via site-directed mutagenesis using 5’p-TCAAGCAGCTCCGGGCAAGAGTCC-3’ (forward) and 5’p-TGCCCCAGACTGTGAGTTGCAACA (reverse) with Platinum SuperFi PCR mastermix (ThermoFisher) as described in the manufacturer’s protocol. The PCR products were circularized using the Phusion T4 Ligase and Rapid Ligation buffer according to the Phusion site-directed mutagenesis kit protocol. A molecular clone containing both the Q577R and L543Q gp41 mutations was constructed by site-directed mutagenesis of pCMV-CK15 (Shingai et al., 2012) using the Phusion site-directed mutagenesis kit (ThermoFisher) as described in the manufacturer’s protocol (for the Q577R substitution) and Platinum SuperFi mastermix (ThermoFisher) (for the L543Q substitution). The Q577R mutation was introduced using the previously described primers, and the L543Q mutation was introduced using 5’p-GGCCAGACAATTATTGTCTGGTAT-3’ (forward) and 5’P-TGTACCGTCAGCGTTATTGACGCT-3’ (reverse) primers. PCR products were circularized using the Phusion T4 Ligase and Rapid Ligation buffer according to the Phusion site-directed mutagenesis kit protocol. 2 μL of the ligation was transformed into TOP10 chemically competent *E. coli* (ThermoFisher), colonies cultured and plasmid DNA extracted as described in the previous section. The mutant virus stocks were prepared as previously described (Shingai et al., 2012).

### Virus replication assay in rhesus monkey PBMC

The preparation and infection of rhesus monkey PBMC have been described previously (Imamichi et al., 2002). Briefly, macaque PBMC, stimulated with concanavalin A and cultured in the presence of recombinant human interleukin-2 (IL-2), were spinoculated (1,200 × *g* for 1 h) with virus at the desired TCID50. Virus replication was assessed by RT assay of the culture supernatant as described above.

### *In vitro* blocking assays with CPT31 in rhesus PBMC

The *in vitro* potency of CPT31 were assessed using a 21-day PBMC replication assay (Nishimura et al., 2002). Briefly, CPT31 was serially diluted (four-fold, starting at 400 nM), and an aliquot of each CPT31 dilution was added to activated rhesus PBMC (1×10^5^ cells per well) in quadruplicate. PBMCs were incubated for 15 minutes and then infected with 100 TCID_50_ of wild-type SHIVAD8-EO or CPT31-resistant SHIVAD8-EO variants. Infected cultures were maintained for 2 weeks, and virus replication was monitored by ^32^P-reverse transcriptase assays (Willey et al., 1988). Any infectious SHIV generated during the 2 weeks of incubation in PBMC would be amplified to levels detectable by this assay.

### CPT31 breadth in vitro

NIH Reagent Panel: The inhibitory potency of CPT31 was tested against a diverse international panel of 60 primary replication competent HIV-1 isolates obtained from the NIH AIDS Reagent Program (#11412) (Brown et al., 2005). Viruses were used as provided without further propagation. Infectious titers were determined using TZM-bl reporter cells (obtained from the NIH AIDS Reagent Program, [#8129]). Cells were grown to ~90% confluency in a 96-well plate prior to addition of serially diluted virus to achieve luminescence signals of 30,000-1,000,000 (BMG Labtech PolarStar Optima plate reader at maximum gain). For low-titer isolates, undiluted virus was used (up to a maximum of 10% of the media volume). Briefly, media with 10 nM, 1 nM, or no CPT31 (uninhibited control) was added to the cells. Virus was subsequently added (up to 10% viral supernatant by volume) to achieve uninhibited luminescence signals of >30,000 (BMG Labtech PolarStar Optima plate reader at maximum gain) and incubated at 37°C for 30 hours. The medium was then removed, cells were lysed using Glo Lysis buffer (Promega), and luciferase substrate (Bright-Glo, Promega) was added as previously described (Welch et al., 2007). Normalized luminescence values were determined by subtracting background luminescence values (TZM-bl cells with no virus) and dividing by the background-subtracted uninhibited control (1.0 = uninhibited, 0 = fully inhibited). Three viral strains from this panel were excluded due to insufficient signal (<30,000) in this assay (57128, TZBD9/11, and E13613M4).

The 118-strain CAVD (Colaboratory for AIDS Vaccine Discovery) pseudoviron panel (Huang et al., 2016; Seaman et al., 2010) was used to measure inhibitor breadth using TZM-BL indicator cells as described.

### CPT31 plasma concentration measurements

CPT31 plasma levels were measured using an LC-MS bioanalytical assay similar to the protocol previously described (Redman et al., 2018). All measurements were made using an Agilent Infinity 1290 HPLC and Agilent 6450A Q-TOF MS with dual Jet Spray source. 200 μL samples were taken from flash-frozen plasma samples (stored at −80°C). Plasma proteins were precipitated by addition of 500 μL 2% NH_4_OH in acetonitrile. After centrifugation to remove precipitated plasma proteins, 500-650 μL of supernatant was transferred to a Waters Oasis MAX 96-well plate (mixed-mode strong anion exchange), washed with 500 μL 2% NH_4_OH in water followed by 500 μL methanol. CPT31 was eluted using 2 × 25 μL 2% formic acid in methanol (for PrEP samples) or 2 × 25 μL or 50 μL 6% formic acid in methanol for the ART rebound samples and treatment samples, respectively.

1-3 μL of this sample was loaded onto an Agilent Accucore 150 C4 HPLC column (2.1 × 50 mm, 2.6 μm pore size). CPT31 was eluted using a gradient between buffer A (20 mM ammonium bicarbonate pH 7.9 in water) and buffer B (acetonitrile). The column was run at 40°C at 0.45 mL/min. Plasma samples were spiked with 5-25 nM of an internal standard CPT31-IS (heavy version of CPT31 with Gly appended to the N-terminus of each of the three PIE12 D-peptides in CPT31, +171.1 Da compared to CPT31). CPT31 and CPT31-IS were monitored using their +6 ions (m/z 1508.5791 and 1537.0899, respectively). The lower limit of quantitation using this method was ~1 nM. Samples were analyzed against an 8-point standard curve of CPT31 fit to a quadratic equation. All data were normalized to the IS, except for samples from the ART rebound study, in which significant IS suppression was observed at high CPT31 concentrations, and the IS was not required to generate a high-quality standard curve.

## Acknowledgements

We thank A. Peach, P. King and B. Yankulova for determining plasma viral RNA loads and R. Petros, C. Ramera, A Bruce and E. Sanford for diligently assisting in the maintenance of animals and assisting with procedures. We thank P. Bjorkman, J. Keeffe, P. Gnanapragasam, A. West, and S. Apple for CAVD panel setup and interpretation and D. Eckert for critical review of the manuscript. We are indebted to Gilead Sciences for providing tenofovir and emtricitabine. We thank the NIH AIDS Research and Reference Reagent Program for TZM-bl cells. This work was supported by the Intramural Research Program of the National Institute of Allergy and Infectious Diseases (MAM), NIH, NIH grants AI076168 and AI150464 (to MSK) and AI95172 (to BDW), and the Bill and Melinda Gates Foundation Collaboration for AIDS Vaccine Discovery (CAVD) grant #OPP1146996 to MSS.

---

Figure S1: **Pharmacokinetic measurements of CPT31 levels during virus acquisition blocking studies.** Four monkeys were intramuscularly administered 3.0 (A), 0.5 (B), or 0.125 (C) mg/kg CPT31 daily, beginning on days –3 to day +7, at each time of SHIVAD8-EO challenge, respectively. Drug levels were measured using an LC-MS bioanalytical assay with an internal standard. The dosing periods are indicated in yellow.

Figure S2: **Pharmacokinetic measurements of CPT31 during monotherapy of chronically infected SHIVAD8-EO rhesus macaques**. CPT31 was administered at 3 mg/kg/day. Drug levels were measured using an LC-MS bioanalytical assay with an internal standard. The dosing period is indicated in yellow.

Figure S3: **Pharmacokinetic measurements of CPT31 in conjunction with cART treatment/cessation in chronically SHIVAD8-EO infected rhesus macaques**. CPT31 was administered at 3 mg/kg/day for 12 weeks between weeks 46 and 58 PI. Drug levels were measured using an LC-MS bioanalytical assay with an internal standard. Dosing period indicated in yellow.

Figure S4: **Sequence analyses of gp41 gene segments in SHIVAD8-EO chronically infected macaques treated with cART and then with CPT31 monotherapy.** Viral RNA was amplified by RT-PCR from plasma collected at week 62. The Q577R changes identified in found in animal DFTV are highlighted.

