## Supplemental Figures and Tables for "Prevention and Treatment of SHIVAD8 Infection in Rhesus Macaques by a Potent D-peptide HIV Entry Inhibitor"

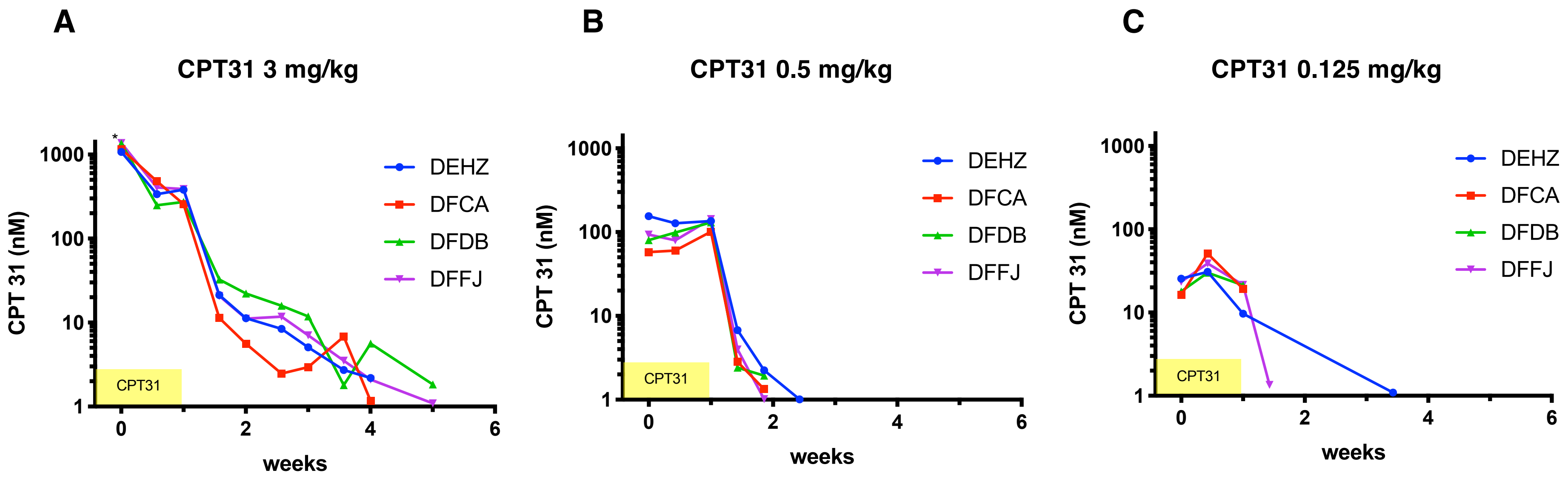


**Figure S1: Pharmacokinetic measurements of CPT31 levels during virus acquisition blocking studies.** Four monkeys were intramuscularly administered 3.0 (A), 0.5 (B), or 0.125 (C) mg/kg CPT31 daily, beginning on days –3 to day +7, at each time of SHIVAD8-EO challenge, respectively. Drug levels were measured using an LC-MS bioanalytical assay with an internal standard. The dosing periods are indicated in yellow.

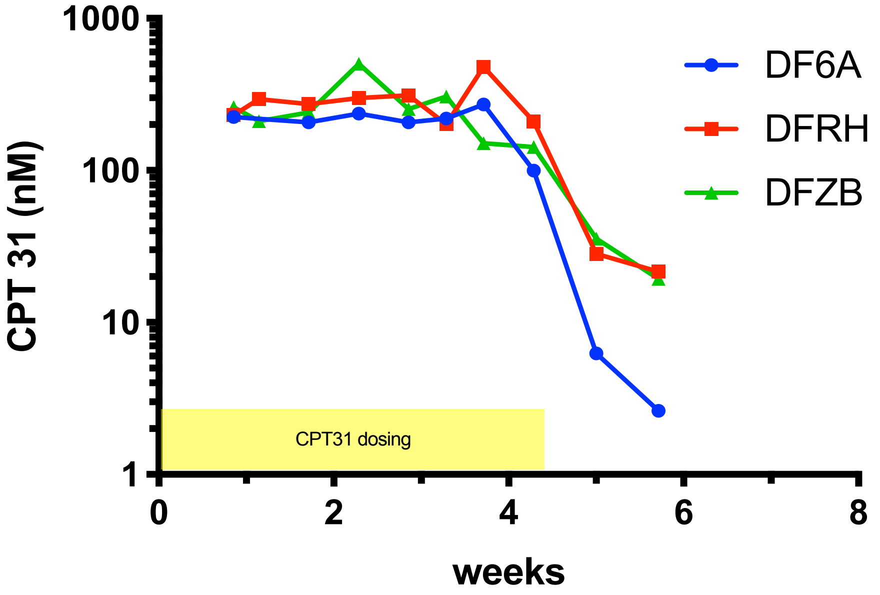


**Figure S2: Pharmacokinetic measurements of CPT31 during monotherapy of chronically infected SHIVAD8-EO rhesus macaques.** CPT31 was administered at 3 mg/kg/day. Drug levels were measured using an LC-MS bioanalytical assay with an internal standard. The dosing period is indicated in yellow.

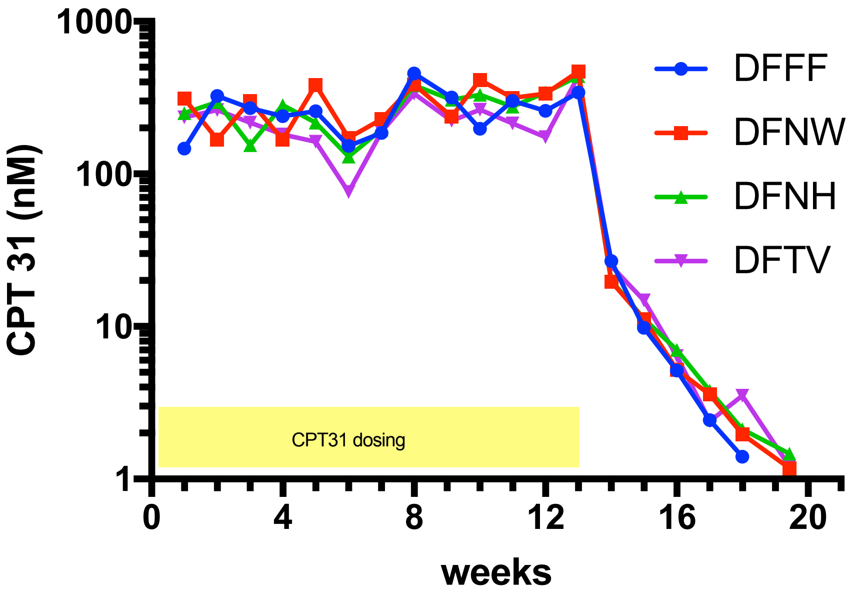


**Figure S3: Pharmacokinetic measurements of CPT31 in conjunction with cART treatment/cessation in chronically SHIVAD8-EO infected rhesus macaques.** CPT31 was administered at 3 mg/kg/day for 12 weeks between weeks 46 and 58 PI. Drug levels were measured using an LC-MS bioanalytical assay with an internal standard. Dosing period indicated in yellow.

```
                                                                                   Q577R

                AVGAIGAMFL GFLGAAGSTM GAASITLTVQ ARLLLSGIVQ QQNNLLKAIE AQQHLLQLTV WGIKQLQARV LAVERYLRDQ QLLGIWGCSG KLICTTAVPW
DFFF (12/21) ---------- ---------- ---------- ---------- ---------- ---------- ---------- ---------- ---------- ----------
DFFF  (3/21) ---------- ---------- ---------- ---------- ------R--- ---------- ---------- ---------- ---------- ----------
DFFF  (2/21) ---------- ---------- ---------- ---------- ---------- ---------- ---------- ---------- ---------- ----------
DFFF  (2/21) ---------- ---------- ---------- ---------- ---------- ---------- ---------- ---------- ---------- ----------
DFFF  (1/21) ---------- ---------- ---------- ---------- ---------- ---------- ---------- ---------- ---------- ----------
DFFF  (1/21) ---------- ---V------ ---------- ---------- ---------- ---------- ---------- ---------- ---------- ----------

DFNH (11/11) -------V-- ---------- ---------- ---------- ---------- ---------- ---------- ---------- ---------- ----------

DFNW (19/30) ---------- ---------- ---------- ---------- ---------- ---------- ---------- ---------- ---------- ----------
DFNW  (5/30) -----R---- ---------- ---------- ---------- ---------- ---------- ---------- ---------- ---------- ----------
DFNW  (3/30) ---------- ---------- ---------- ---------- ---------- ---------- ---------- ---------- ---------- ----------
DFNW  (3/30) ---------- ---------- ---------- ---------- ---------- ---------- ---------- ---------- ---------- ----------

DFTV (12/21) ---------- ---------- ---------- ---------- ---------- ---------- ---------- ---------- ---------- ----------
DFTV  (7/21) ---------- ---------- ---------- --Q------- ---------- ---------- ------R--- ---------- ---------- ----------
DFTV  (2/21) ---------- ---------- ---------- ---------- ---------- ---------- ------R--- ---------- ---------- ----------


                NASWSNKTLD MIWNNMTWME WEREIDNYTG LIYTLIEESQ NQQEKNEQEL LELDKWASLW NWFDITNWLW YIK
DFFF (12/21) ---------- ---------- ---------- ---------- ---------- ---------- ---------- ---
DFFF  (3/21) ---------- ---------- ---------- ---------- ---------- ---------- ---------- ---
DFFF  (2/21) ---------- ---------- -----N---- ---------- ---------- ---------- ---------- ---
DFFF  (2/21) ---------- ---------- --G------- ---------- ---------- ---------- ---------- ---
DFFF  (1/21) ---------- ---------- ---------- -----L---- ---------- -------G-- ---------- ---
DFFF  (1/21) ---------- ---------- ---------- ---------- ---------- ---------- ---------- ---

DFNH (11/11) ---------- ---------- ---------- ---------- ---------- ---------- ---------- ---

DFNW (19/30) ---------- ---------- ---------- ---------- ---------- ---------- ---------- ---
DFNW  (5/30) ---------- ---------- ---------- ---------- ---------- -------G-- ---------- ---
DFNW  (3/30) ---------- ---------- ------D--D ---------- ---------- ---------- ---------- ---
DFNW  (3/30) ---------- ---------- ---------D ---------- ---------- ---------- ---------- ---

DFTV (12/21) ---------- ---------- ---------- ---------- ---------- ---------- ---------- ---
DFTV  (7/21) ---------- ---------- ---------- ---------- ---------- ---------- ---------- ---
DFTV  (2/21) ---------- ---------- ---------- ---------- ---------- ---------- ---------- ---
```

**Figure S4: Sequence analyses of gp41 gene segments in SHIVAD8-EO chronically infected macaques treated with cART and then with CPT31 monotherapy.** Viral RNA was amplified by RT-PCR from plasma collected at week 62. The Q577R changes identified in found in animal DFTV are highlighted.

Table S1: Inhibitory breadth of CPT31 against the CAVD 118-strain pseudovirion panel.

| Strain | Clade | CPT31 $IC_{50}$ (pM) |
|---|---|---|
| 191084_B7_19 | A | 1 |
| 211_9 | A | 2 |
| 928_28 | A | 3 |
| 0815_V3_C3 | A | 4 |
| 3365_V2_C20 | A | 4 |
| MS208_A1 | A | 10 |
| 6540_V4_C1 | A | 10 |
| Q23_17 | A | 20 |
| Q461_E2 | A | 20 |
| 191955_A11 | A | 20 |
| T251_18 | A | 20 |
| 9004SS_A3_4 | A | 20 |
| Q769_D22 | A | 20 |
| 235_47 | A | 30 |
| T278_50 | A | 30 |
| 263_8 | A | 30 |
| 3817_V2_C59 | A | 30 |
| 6041_V3_C23 | A | 40 |
| 6545_V4_C1 | A | 40 |
| Q842_D12 | A | 50 |
| T250_4 | A | 60 |
| 3415_V1_C1 | A | 60 |
| T255_34 | A | 60 |
| 6480_V4_C25 | A | 80 |
| Q259_17 | A | 80 |
| 0260_V5_C36 | A | 200 |
| T257_31 | A | 240 |
| R2184_C4 | AE | 3 |
| C3347_C11 | AE | 20 |
| CNE8 | AE | 20 |
| C2101_C1 | AE | 20 |
| R3265_C6 | AE | 20 |
| R1166_C1 | AE | 20 |
| CNE5 | AE | 30 |
| BJOX028000_10_3 | AE | 30 |

| | | |
|---|---|---|
| C1080_C3 | AE | 40 |
| 620345_C1 | AE | 50 |
| BJOX025000_01_1 | AE | 50 |
| C4118_9 | AE | 80 |
| BJOX010000_06_2 | AE | 90 |
| BJOX009000_02_4 | AE | 160 |
| BJOX015000_11_5 | AE | 190 |
| AC10_29 | B | 10 |
| SC422661_8 | B | 10 |
| REJO4541_67 | B | 20 |
| SC05_8C11_2344 | B | 20 |
| 62357_14_D3_4589 | B | 20 |
| 1012_11_TC21_3257 | B | 20 |
| 1054_07_TC4_1499 | B | 30 |
| WEAU_D15_410 | B | 30 |
| WITO4160_33 | B | 30 |
| TRJO4551_58 | B | 30 |
| TRO_11 | B | 40 |
| 6535_3 | B | 40 |
| THRO4156 | B | 50 |
| CAAN5342 | B | 60 |
| 1056_10_TA11_1826 | B | 60 |
| RHPA4259_7 | B | 70 |
| 6240_08_TA5_4622 | B | 80 |
| 6244_13_B5_4567 | B | 100 |
| QH0692 | B | 110 |
| 1006_11_C3_1601 | B | 140 |
| PVO_4 | B | 220 |
| ZM233_6 | C | 4 |
| DU422 | C | 10 |
| 249M_B10 | C | 10 |
| DU156_12 | C | 10 |
| DU172_17 | C | 10 |
| 0013095_2_11 | C | 10 |
| CE0393_C3 | C | 10 |
| ZM249_1 | C | 10 |
| ZM135_10A | C | 10 |
| CE704809221_1B3 | C | 20 |
| CNE58 | C | 20 |

| | | |
|---|---|---|
| CNE53 | C | 20 |
| ZM53_12 | C | 20 |
| CAP210_E8 | C | 20 |
| ZM214_15 | C | 20 |
| 6811_V7_C18 | C | 20 |
| CNE20 | C | 20 |
| ZM109_4 | C | 20 |
| CE1086_B2 | C | 20 |
| CNE17 | C | 20 |
| CNE19 | C | 20 |
| ZM197_7 | C | 20 |
| CE1172_H1 | C | 20 |
| CNE52 | C | 30 |
| 16055_2_3 | C | 30 |
| CNE21 | C | 30 |
| CAP45_G3 | C | 30 |
| CE2060_G9 | C | 30 |
| ZM247V1(REV_) | C | 30 |
| CE0682_E4 | C | 30 |
| 3301_V1_C24 | C | 40 |
| 246F_C1G | C | 40 |
| 001428_2_42 | C | 40 |
| 1394C9G1(REV_) | C | 60 |
| CE703010054_2A2 | C | 60 |
| 3103_V3_C10 | C | 70 |
| BF1266_431A | C | 80 |
| CNE30 | C | 80 |
| 16845_2_22 | C | 80 |
| CE1176_A3 | C | 110 |
| 7030102001E5(REV_) | C | 190 |
| CE2010_F5 | C | 490 |
| 231966_C2 | D | 10 |
| 6952_V1_C20 | D | 20 |
| A07412M1_VRC12 | D | 50 |
| 89_F1_2_25 | D | 60 |
| 3016_V5_C45 | D | 80 |
| 6405_V4_C34 | D | 140 |
| 231965_C1 | D | 240 |
| X1254_C3 | G | 10 |

| | | |
|---|---|---|
| X1193_C1 | G | 20 |
| X1632_S2_B10 | G | 20 |
| X2088_9 | G | 20 |
| P1981_C5_3 | G | 30 |
| X2131_C1_B5 | G | 30 |
| P0402_C2_11 | G | 40 |

Table S2: Inhibitory breadth of CPT31 against 60-strain international panel of replication-competent virus obtained from the NIH AIDS Reagent Program. Note that three strains were excluded due to insufficient titer (see Methods).

| Strain | Clade | Normalized luminescence | |
|---|---|---|---|
| | | 1 nM CPT31 | 10 nM CPT31 |
| 92UG029 | A | 0.048 | 0.018 |
| KER2008 | A | 0.035 | 0.011 |
| KER2018 | A | 0.052 | 0.025 |
| KNH1088 | A | 0.015 | 0.004 |
| KNH1135 | A | 0.046 | 0.024 |
| KNH1144 | A | 0.079 | 0.015 |
| KNH1207 | A | 0.079 | 0.018 |
| KNH1209 | A | 0.057 | 0.048 |
| KSM4030 | A | 0.064 | 0.044 |
| 93RW024 | A | 0.03 | 0.007 |
| M02138 | AE | 0.048 | 0.02 |
| CM235/GS020 | AE | 0.162 | 0.017 |
| CM244 | AE | 0.046 | 0.005 |
| CM240/GS022 | AE | 0.134 | 0.024 |
| NI1046 | AE | 0.009 | 0.003 |
| NI1052 | AE | 0.02 | 0.003 |
| NI1149 | AE | 0.071 | 0.026 |
| NP1251 | AE | 0.03 | 0.016 |
| NP1525 | AE | 0.012 | 0.002 |
| NP1695 | AE | 0.043 | 0.03 |
| 55815 | AG | 0.012 | 0.005 |
| CAM0002 | AG | 0.044 | 0.003 |
| CAM0013 | AG | 0.016 | 0.003 |
| CAM0014 | AG | 0.012 | 0.005 |
| CAM0015 | AG | 0.04 | 0.03 |
| CAM0005 | AG | 0.016 | 0.002 |
| CAM0008 | AG | 0.032 | 0.003 |
| CAM1475MV | AG | 0.017 | 0.012 |
| CAM1970LE | AG | 0.027 | 0.015 |
| DJ263/GS003 | AG | 0.033 | 0.008 |
| 873 | B | 0.064 | 0.013 |
| 33931N | B | 0.024 | 0.009 |
| Ba-L | B | 0.031 | 0.002 |
| BK132/GS009 | B | 0.003 | 0.002 |
| BX08 | B | 0.058 | 0.005 |
| BZ167 | B | 0.006 | 0.002 |
| MN/H9 | B | 0.007 | 0 |
| NP1538 | B | 0.05 | 0.006 |
| US1/GS0004 | B | 0.047 | 0.023 |
| US4/GS007 | B | 0.089 | 0.013 |
| 56313 | C | 0.021 | 0.007 |
| 20635-4 | C | 0.01 | 0.004 |
| PBL286 | C | 0.042 | 0.01 |
| PBL288 | C | 0.97 | 0.88 |

| | | | |
|---|---|---|---|
| SE364/GS015 | C | 0.084 | 0.023 |
| SM145/GS016 | C | 0.694 | 0.243 |
| TZA246 | C | 0.017 | 0.016 |
| TZA68 | C | 0.023 | 0.003 |
| 301965 | C | 0.039 | 0.023 |
| 93UG065 | D | 0.063 | 0.056 |
| A03349M1 | D | 0.064 | 0.005 |
| A07412M1 | D | 0.08 | 0.033 |
| A08483M1 | D | 0.062 | 0.026 |
| D26830M4 | D | 0.047 | 0.015 |
| E08464M4 | D | 0.068 | 0.019 |
| J32228M4 | D | 0.039 | 0.02 |
| NKU3006 | D | 0.028 | 0.002 |
